# Signs of host genetic regulation in the microbiome composition in cattle

**DOI:** 10.1101/100966

**Authors:** O. Gonzalez-Recio, I. Zubiria, A. García-Rodríguez, A. Hurtado, R. Atxaerandio

**Author notes:** Corresponding author; Oscar González-Recio.

## Abstract

Previous studies have revealed certain genetic control by the host over the microbiome composition, although in many species the host genetic link controlling microbial composition is yet unknown. This potential association is important in livestock to study all factors and interactions that rule the effect of the microbiome in complex traits. This report aims to study whether the host genotype exerts any genetic control on the microbiome composition of the rumen in cattle. Data on 16S and 18S rRNA gene-based analysis of the rumen microbiome in 18 dairy cows from two different breeds (Holstein and Brown Swiss) were used. The effect of the genetic background of the animal (through the breed and Single Nucleotide Polymorphisms; SNP) on the relative abundance (RA) of archaea, bacteria and ciliates (with average relative abundance per breed >0.1%) was analysed using Bayesian statistics. In total, 13 genera were analysed for bacteria (5), archaea (1), and ciliates (7). All these bacteria and archaea genera showed association to the host genetic background both for breed and SNP markers, except RA for the genera *Butyrivibrio* and *Ruminococcus* that showed association with the SNP markers but not with the breed composition. Relative abundance of 57% (4/7) of ciliate analysed showed to be associated to the genetic background of the host. This host genetic link was observed in some genus of *Trichostomatia* family. For instance, the breed had a significant effect on *Isotricha*, *Ophryoscolex* and *Polyplastron*, and the SNP markers on *Entodinium*, *Ophryoscolex* and *Polyplastron*. In total, 77% (10/13) of microbes analysed showed to be associated to the host genetic background (either by breed or SNP genotypes). Further, the results showed a significant association between DGAT1, ACSF3, AGPAT3 and STC2 genes with the relative abundance *Prevotella* genus with a false discovery rate lower than 15%. The results in this study support the hypothesis and provide some evidence that there exist a host genetic component in cattle that can partially regulate the composition of the microbiome.

---

The research interest on the microbiome and its effects on the host, both in humans [1,2] and livestock [3,4], is raising in the last years. The microbiome plays an important role in the phenotypic expression of many phenotypes such as feed efficiency, disease status, or methane emission. Traditionally, microbes have been studied in the lab, without considering their effect on complex features and their interaction with the host. In the particular case of livestock, the traits of interest are usually related to productive, health or environmental factors. In the last decade, more attention has been focused on the interactions between microbes and diet [5–8], methane emissions [9–13] and the microbiome compositions across hosts, environment and age [4,7,14]. It has also been proposed as a predictor of complex traits [13,15].

Therefore, there is an increasing interest on determining whether a host genetic control on the microbiome composition exists. Recent studies show some evidences that support the hypothesis that there is some sort of host control over the composition of the microbiome in mammals. For instance, Weimer et al. (2010) reported that after a near-total exchange of ruminal contents, the ruminal bacterial composition returned to a similar status to that prior the exchange. More recently, [17] showed differences between sire progeny groups on the archaea:bacteria ratio in Aberdeen Angus and Limousin cattle breeds, and [18] reported heritabilities above 0.20 for the relative abundance of several microbes in a twin human study.

It is of interest to provide more evidences on the host genetic control of the microbiome composition because some selection intensity could be applied to select individuals with a favourable microbiome for a given breeding goal, as the reduction of methane yield or the improvement of the feed efficiency, for example.

This trial was carried out in accordance with Spanish Royal Decree 53/2013 for the protection of animals used for experimental and other scientific purposes. In this study, ruminal content was sampled from 18 dairy cows (10 Holstein and 8 Brown Swiss) from Fraisoro Farm School (Zizurkil, Gipuzkoa, Spain). These cows were undergoing a nutrition experiment. They were randomly assigned to one of two experimental concentrate supplements. Concentrates were formulated to contain cold-pressed rapeseed cake or palm as fat sources, and to provide equal amounts of crude protein, energy and fat. Both breeds were fed both diets. The effect of the treatment was adjusted as a 2-levels factor in the statistical analyses, but results are not reported here as this is not the objective of this study.

Rumen samples were taken 4 times over two consecutive days. Sampling began at 00:00 and 12:00 h on d 1, and 06:00 and 18:00 h on d 2; each sampling taking approximately 2 h. Ruminal samples were collected from each dairy cow using a stomach tube connected to a mechanical pumping unit. About 100 ml of each ruminal extraction were placed into a container and were frozen immediately after the extraction and then stored at −20±5°C until analysis.

Then, samples were gradually thawed overnight at refrigeration (5±3°C) and squeezed through four layers of sterile cheesecloth to separate solid (solids with a particle size smaller than the diameter of the tube) from liquid digesta phases. This latter phase was subsequenty separated into planktonic organisms and bacteria associated with the liquid fraction. The solid phase was separated in associated and adherent fractions. Fractionation procedures were carried out following the methodology described in [19]. The four fractions were lyophilized and composited to obtain a unique sample with the four fractions represented proportionally (on dry matter basis).

After composition, DNA extraction was performed using the commercial Power Soil DNA Isolation kit (Mo Bio Laboratories Inc, Carlsbad, CA, USA) following manufacturer’s instructions. The extracted DNA was subjected to paired-end Illumina sequencing of the V4 hypervariable region of the 16S rRNA [20] and of the V7 region of the 18S rRNA genes. The libraries were generated by means of Nextera kit. The 250 bp paired-end sequencing reactions were performed on a MiSeq platform (Illumina, San Diego, CA, USA).

Sequence data were processed using the QIIME software package version 1.9.1 [21]. Sequences below 220 bp in length and Phred score below 20 were discarded. In total, 3,261,168 and 3,431,242 reads from the 16S and 18S rRNA regions respectively, were analysed. Sequence data were grouped into operational taxonomic units (OTU) sharing 97% sequence similarity, and assigned to phylogenetic groups by BLAST [22].

Bacterial and archaeal 16S rRNA genes were assigned using the GreenGenes database (May 2013 version) and ciliate protozoal 18S rRNA genes against SILVA database (March 2015 version). Data were summarised at the genus level. Relative abundance (RA) of genera in each animal was calculated after excluding those genera that appeared in <0.1% proportion in the previous step. Only genera showing average RA>0.1% in both breeds were kept for subsequent analyses.

Genotypes from animals under study were also obtained with the Illumina 9K chip (Illumina, Inc, San Diego, CA, USA). A total of 9,146 SNPs with minor allele frequency (MAF) >0.05 in the whole genotyped Spanish population were kept (data provided by the Spanish Holstein association www.conafe.com from more than 3,000 individuals).

We used two strategies to analyse the host genetic effect on the microbiome composition. Our response variable was the RA of the most common ruminal microbes previously found, and the model adjusted by diet treatment (2 groups, with or without cold-pressed rapeseed cake) and age (primiparous *vs* multiparous) groups and days in milk as a covariate. In the first strategy, differences at the breed level (Holstein *vs* Brown Swiss) were estimated (Model 1). Model 1)

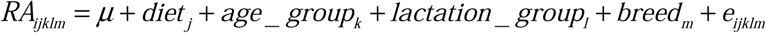

The second strategy included the first two principal components (PC) of a genomic relationship matrix instead of the breed effect as model 2)

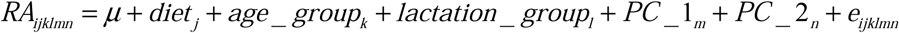

This genomic relationship matrix was calculated as in [23], where the genome relationship between individuals *i* and *j* can be calculated as

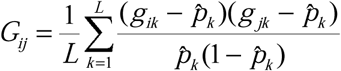

where *g_ik_* refers to the gene frequency value genotypes *AA*, *Aa* and *aa*, coded as 1, 0.5 and 0, respectively, of individual *i* or *j* at locus *k* (*k* = 1, 9146). Gene frequency is half the number of copies of the reference allele *A*. Then, *p̂_k_* was the estimated allele frequency in the whole genotyped population as provided by CONAFE. The first PC of this matrix aims to detect stratification at the breed level (Holstein *vs* Brown Swiss), whereas the subsequent PC are expected to capture genomic differences between individuals.

Bayesian analyses were performed to estimate the breed and principal component effects [24] using an in-home suite of programs written in R software [25]. Evidence of a host genetic effect was considered when the 80% of the posterior distribution for the breed or the PC had the same sign (either positive or negative). This is, 80% of the posterior probability for the respective effect fell either above or below zero. Those microorganisms that showed evidence of a host genetic control were selected to implement genome wide association analyses. Here, the RA of those microorganisms was used as a dependent variable, and the SNP markers were selected as explicative variables in a single marker linear regression model, including breed and diet as environmental factors. The p-values were adjusted on false discovery rate (FDR).

The gene content of the significant SNP was examined using the bovine genome annotation in BioMart tool of Emsembl (ensembl.org/biomart) using Ensembl Genes 75 database. The National Center for Biotechnology Information (NCBI) database and PubMed were employed to investigate the potential biological relation of the genes that contained the SNP and the microbes in order to propose candidate genes that underlie the detected associations.

## RESULTS AND DISCUSSION

The results from the 16S rRNA region showed a 98:2 for the bacteria:archaea ratio. The more abundant bacterial phyla were *Bacteroidetes* (58%), *Firmicutes* (33%) and TM7 *(Candidatus Saccharibacteria)* (4%). Methanobacteria was the most abundant clade among the archaeas. Taxa composition was similar to those reported before in other ruminal microbiome communities [7,13], being mainly microbes related to peptide and cellulose degradation or to the synthesis of microbial protein and volatile fatty acids.

### Bacteria and archaea

The RA of genera analysed are shown in Figure 1. *Prevotella* was the most abundant bacteria-archaea genus in both breeds, followed by *Butyrivibrio* and *Succiniclasticum*. The archaea *Methanobrevibacter* was more abundant than the rest of the archaea genera detected in the samples.

**Figure 1.**
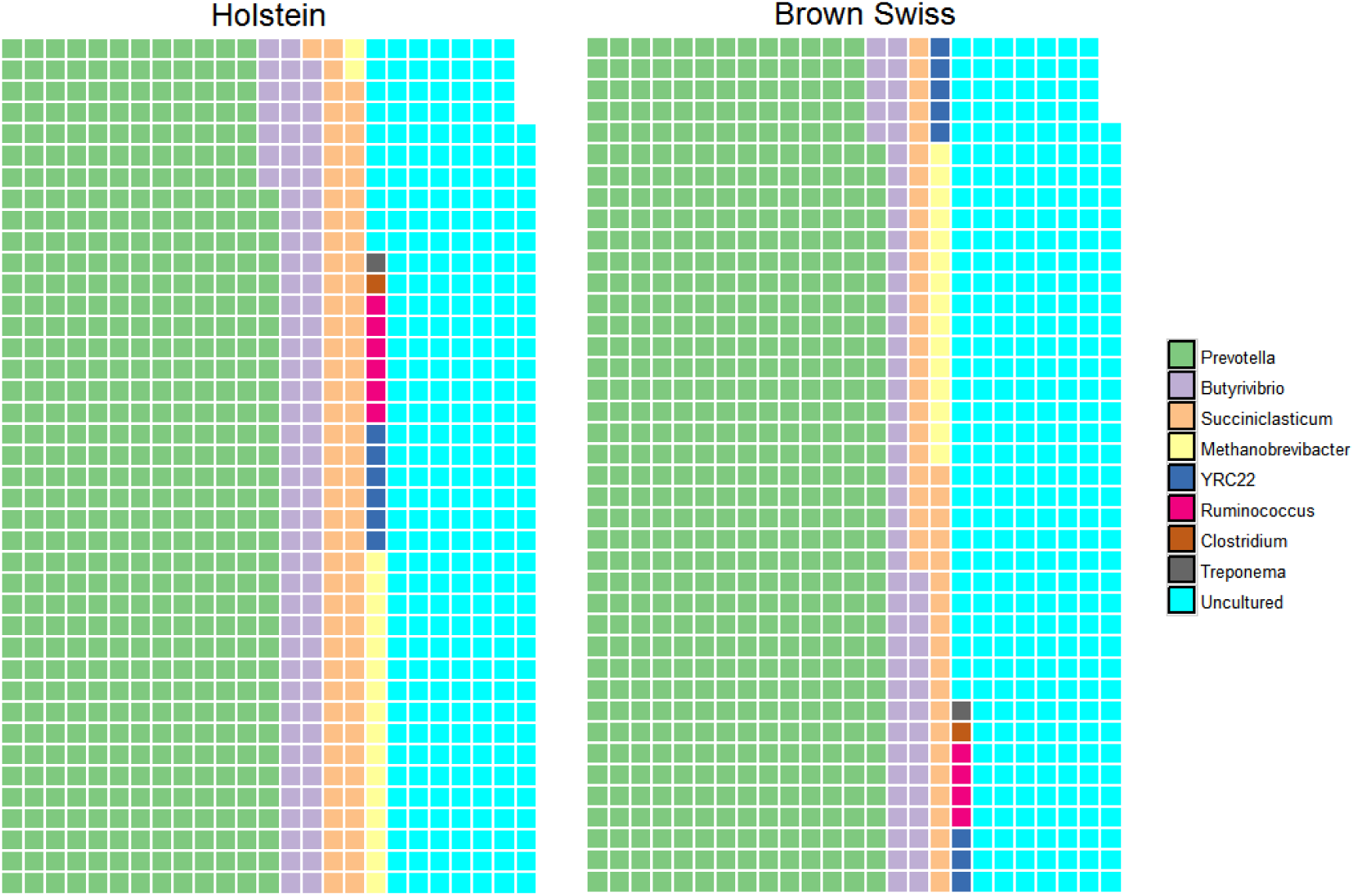
Relative abundance of Bacteria and Euryarchaea with average relative abundance larger than 0.1% in both breeds.

Table 1 shows the results from the statistical analyses of the host genetic component on the different RA. The analyses showed differences between breeds for 4 (*Methanobrevibacter*, *Succiniclasticum*, *Prevotella* and *YRC22*) out of the 6 archaea and bacteria genera analysed from 16S rRNA region. However, either the first or second genomic PC were significant for all other genus analysed (Table 1).

**Table 1.**
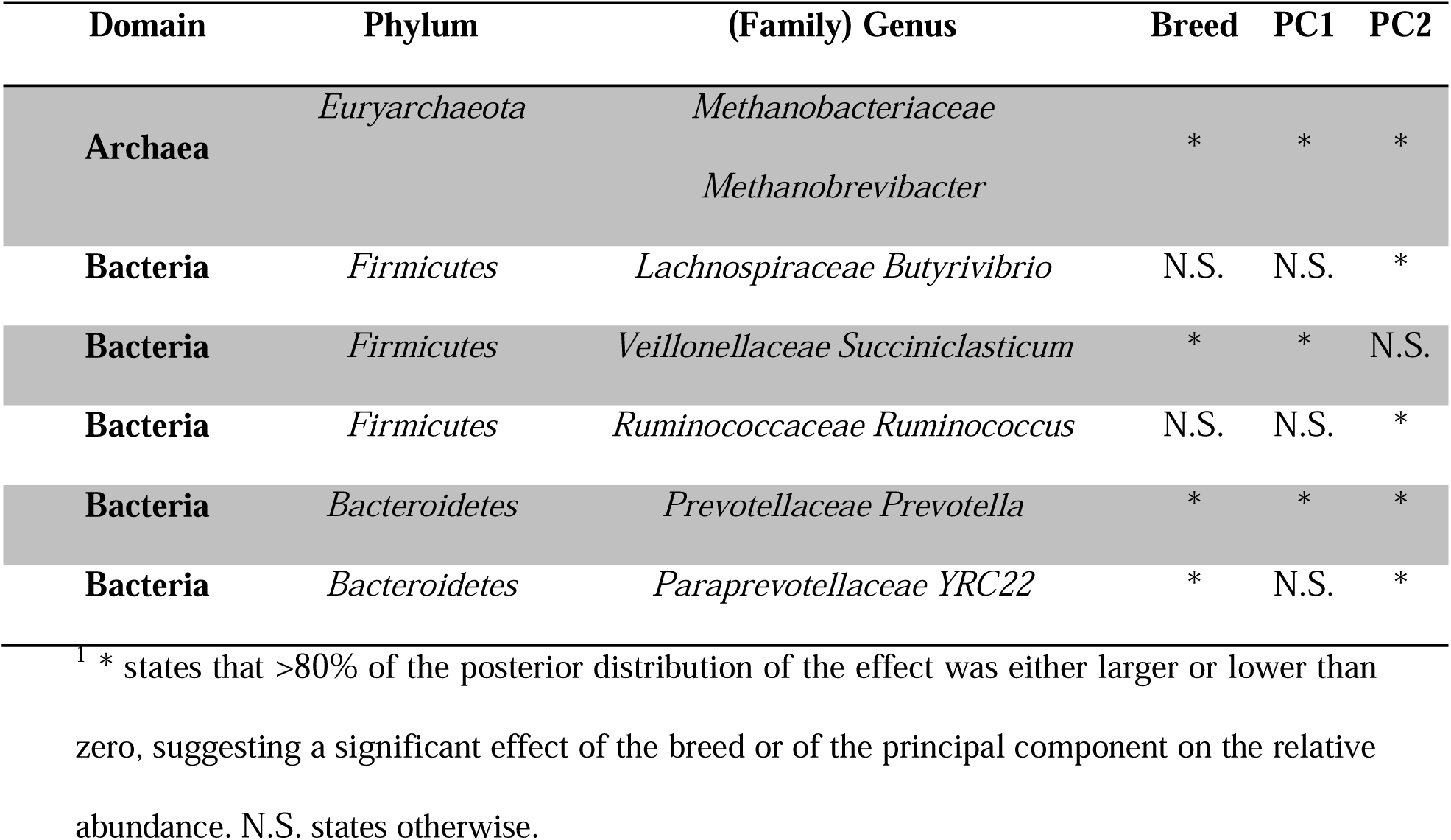
Effect^1^ of the breed (Holstein vs Browm Swiss) and the two first principal component of a genomic relationship matrix based on genotypes on the relative abundance of different bacteria and archaea genera. Only genera that are present with average relative abundance larger than 0.1% in both breeds are shown.

| Domain | Phylum | (Family) Genus | Breed | PC1 | PC2 |
| --- | --- | --- | --- | --- | --- |
| Archaea | Euryarchaeota | Methanobacteriaceae<br>Methanobrevibacter | * | * | * |
| Bacteria | Firmicutes | Lachnospiraceae Butyrivibrio | N.S. | N.S. | * |
| Bacteria | Firmicutes | Veillonellaceae Succiniclasicum | * | * | N.S. |
| Bacteria | Firmicutes | Ruminococcaceae Ruminococcus | N.S. | N.S. | * |
| Bacteria | Bacteroidetes | Prevotellaceae Prevotella | * | * | * |
| Bacteria | Bacteroidetes | Paraprevotellaceae YRC22 | * | N.S. | * |

<sup>1</sup> \* states that >80% of the posterior distribution of the effect was either larger or lower than zero, suggesting a significant effect of the breed or of the principal component on the relative abundance. N.S. states otherwise.

### Ciliate

Figure 2 shows the relative abundance of the analysed ciliate in both breeds. The genus *Entodinium* was the most abundant among the ciliate protozoal, followed by *Isotricha*. Phenotypically, *Ophryoscolex*, *Diplodinium* and *Polypastron* were more abundant in Holstein, whereas *Dasytricha* showed larger RA in Brown Swiss. The breed effects showed differences in 3 (*Isotricha*, *Ophryoscolex* and *Polyplastron*) out of 7 ciliate genus analysed. The genomic PCs were also statistically significant for these genera, except for *Isotricha*, where the posterior distribution did not show a significant effect for the PCs (Table 2).

**Figure 2.**
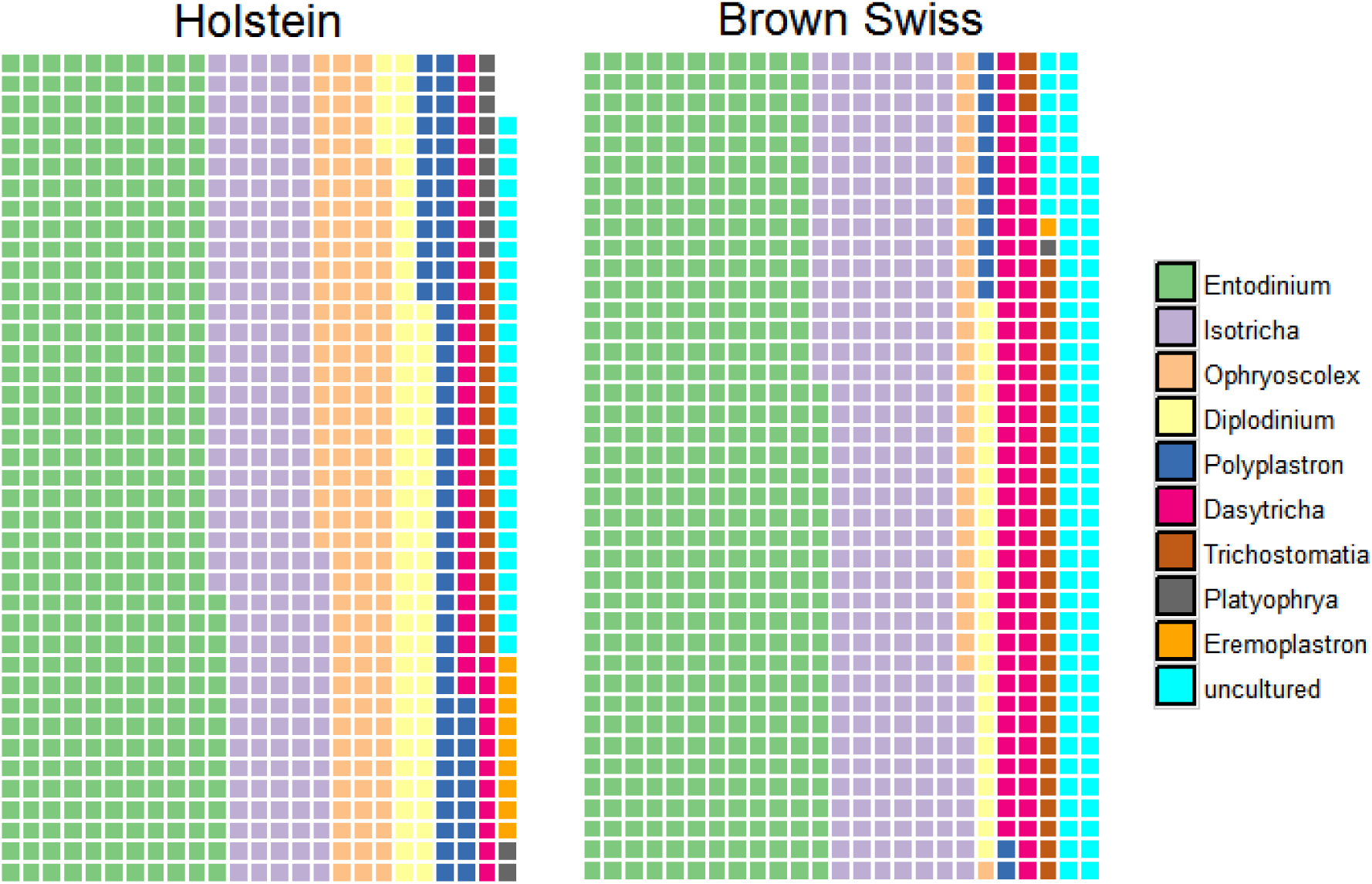
Relative abundance of genera of ciliate with average relative abundance larger than 0.1% in both breeds.

**Table 2.** Effect^1^ of the breed (Holstein vs Brown Swiss) and the two first principal component of a genomic relationship matrix based on genotypes on the relative abundance of different ciliate genera. Only genera that are present with average relative abundance larger than 0.1% in both breeds are shown.

| Domain | Order | (Family) Genus | Breed | PC1 | PC2 |
| --- | --- | --- | --- | --- | --- |
| **Eukaryota** | *Ciliophora* | *Litostomatea Trichostomatia* | N.S. | N.S. | N.S. |
| **Eukaryota** | *Ciliophora* | *Trichostomatia Dasytricha* | N.S. | N.S. | N.S. |
| **Eukaryota** | *Ciliophora* | *Trichostomatia Entodinium* | N.S. | N.S. | * |
| **Eukaryota** | *Ciliophora* | *Trichostomatia Isotricha* | * | N.S. | N.S. |
| **Eukaryota** | *Ciliophora* | *Trichostomatia Diplodinium* | N.S. | N.S. | N.S. |
| **Eukaryota** | *Ciliophora* | *Trichostomatia Ophryoscolex* | * | N.S. | * |
| **Eukaryota** | *Ciliophora* | *Trichostomatia Polyplastron* | * | * | * |

<sup>1</sup> \* states that >80% of the posterior distribution of the effect was either larger or lower than zero, suggesting a significant effect of the breed or of the principal component on the relative abundance. N.S. states otherwise.

Despite the small sample size, RA for 77% (10/13) of the genera analysed were found to be regulated by some host genetic factor (breed, SNP marker, or both), which suggest that the microbiome composition is regulated by some genetic mechanisms in the host. The host genetic background showed to have effect on a larger proportion of bacteria-archaea, in comparison to ciliates. We did not find a host genetic effect on the relative abundance of genera *Trichostomatia*, *Dasytricha*, and *Diplodinium*. These microbes might be more influenced by diet than by the host genetic effect, and larger sample sizes might be neccesary to detect differences between breeds or host genetic effects.

[18] also showed host genetic effect on the RA of different genera and families of *Firmicutes* and *Euryarchaeota* (e.g. *Turicibacter*, *Blautia Clostridiaceae*, *Ruminococcaceae* or *Methanobrevibacter*) in humans. Their study also showed a host genetic effect on some *Tenericutes*, *Proteobacteria* (Family *Oxalobacteraceae*) and *Actinobacteria* (Genera *Bifidobacterium* and *Actinomyces*). Our study also shows a host genetic effect on some genera of Firmicutes but also on some *Bacteroidetes* differently to [18] and ciliate which were not analysed in the human study as they are not abundant in the human gut. Roehe et al. [17] showed differences in the microbial community of progeny daughters from different cattle breeds and sires, suggesting that even under the same diet and environmental circumstances, individuals can differ in their microbial communities depending on their progenitors.

Microbial networks for 16S and 18S-gene rRNA regions were constructed using the algorithm described by [26] and their graphical representations are shown in Figures S1 and S2. The microorganisms that showed to be related to the host genetics are relevant in the composition of the ruminal environment and the degradation of feed. For instance, bacteria from the *Prevotella*, the most abundant group, and *Paraprevotella* genera are involved in the metabolism of proteins and peptides in the rumen. They break down protein and carbohydrates in feed [27], synthesize *de novo* peptides and use products of cellulose degradation from other cellulolytic bacteria [28,29]. Further, bacteria from the genus *Ruminococcus* break down cellulose, hemicellulose and produce succinic acid as a major fermentation product together with acetic and formic acids, hydrogen and CO_2_. These products are then used by other bacteria, some from the *Succiniclasticum* genus, which convert succinate to propionate as an energy-yielding mechanism. *Butyrivibrio* bacteria are proteolytic bacteria and are involved in the degradation of hemicellulose walls, and lipid hydrogenation. They produce mainly butyrate, that is metabolized through the rumen wall to produce energy. Further, archaeas from the *Methanobrevibacter* genus use hydrogen and CO_2_ products and by-products from other microorganism (e.g. *Ruminococcus*) to synthesise methane. The archaea, mainly organisms related with genera *Methanobrevibacter* and *Methanosphaera*, are highly associated with methane emission in ruminants [27].*Methanobrevibacter* has been associated to methane emissions in many previous studies, e.g. [13,30,31].

*Entodinium* ciliate are able to engulf small plant particles and degrade cellulose [27,32].They are considered as cellulolytic microorganisms. *Isotricha* and *Dasytricha* use soluble sugar, and many carbohydrates enzymatic activities have been detected. *Polypastron* ciliates can actively ingest large cellulosic fibres of the rumen fluid [27,33]. The products of rumen ciliates are more or less similar and include acetate, butyrate, and lactate. They also produce CO_2_ and hydrogen during the synthesis that can be converted to methane by methanogenic archaea and protozoa. Ciliates interact with other rumen microorganism as they can ingest bacteria as protein source. A host genetic effect on the RA of these microorganisms explain the heritability found in related traits such as feed efficiency or methane yield [34–36].

Genome-wide association analyses was performed for the RA the four microorganisms that showed significant effect on both breed and PC1 effect (*Methanobrevibacter*, *Succiniclasticum*, *Prevotella*, and *Polyplastron* genera). The generalized linear model implemented included the breed, diet and the bovine SNP marker effects, and p-values were adjusted on false discovery rate (FDR). As expected, the small sample size caused that most of the markers with significant P-values (<0.01) presented a large FDR. We chose the threshold of FDR<0.15 (equivalent P-value of 1.81×10^−4^) to report significant SNP markers. After this adjustment, significant bovine SNP markers were found for *Prevotella* genus RA (Table 3). Most of these markers were within known genes with functions related mainly to metabolic pathways and signalling on the central neural system. The role of the microbiome in the metabolic status and the development of several central system disorders have been well establish in humans [37,38], and our results suggest that there are also associations between genes involved in metabolic and neural processes and the rumen microbiota compositions. It must be highlighted that we found association between the DGAT1 gene and the RA of P. Prevotella. The DGAT1 gene is a major gene with a large effect on the fat composition in milk [39–41]. The association found in this study shows that the effect of the DGAT1 on the milk fat composition may be partially regulated by some effect on the microbiome composition, where individuals carrying the A (A/G) allele of the ARS-BFGL-NGS-4939 SNP tend to host a larger proportion of *Prevotella* microorganisms which are also involved in the protein and peptide degradation in the rumen, in the production of saturated fatty acids as well as in saccharolytic pathways. Other genes with significant association to the *Prevotella* RA were the ACSF3, AGPAT3 and STC2, all of them previously associated to fatty acids or cell metabolism.

**Table 3.**
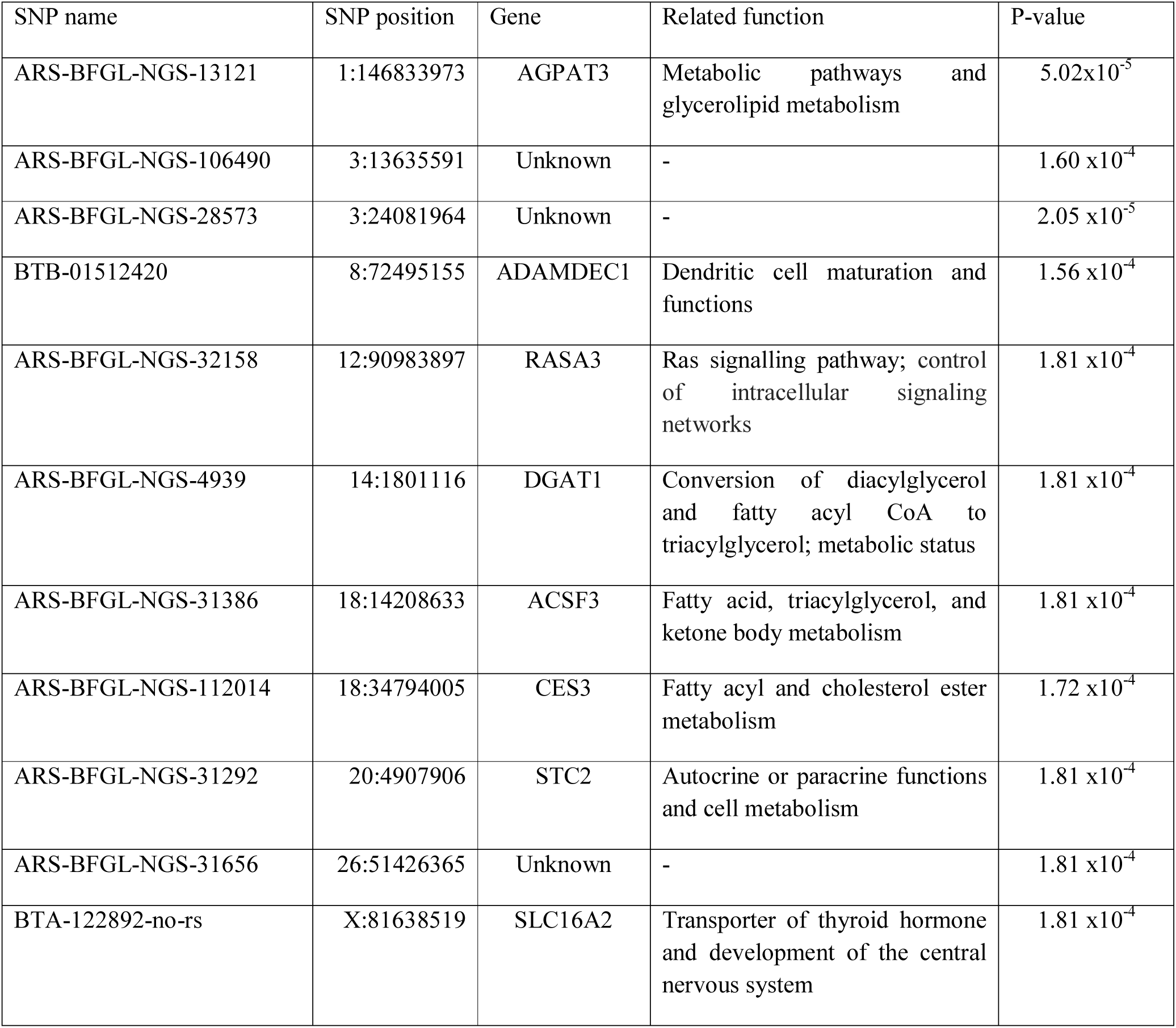
*Genes contained within significant bovine SNP markers for the relative abundance of P. Prevotella, and their position*.

The results in this study provide some evidence that support the hypothesis of a host genetic component that can partially regulate the composition of the microbiome, and indirectly some metabolic pathways. In this sense, it seems that there is a genetic component in the regulation of some groups of H2- producing microorganisms included in the *Firmicutes* phylum and ciliate protozoa and H_2_-utilizers bacteria associated to *Bacteroidetes*. This is relevant because diets and management practices can be specifically designed to compensate those genotypes that are more susceptible to harbour less efficient microorganisms from a nutritional and energetic point of view. Results from this study must be considered carefully due to the reduced sample size. Future studies should allow to better estimate heritability of the microbiome composition in cattle, as well as covariance components with other traits of interest (e.g. feed efficiency, productivity, or methane emissions). Still, if these results were confirmed, breeding strategies could be developed to select future livestock generations prone to harbor a favourable microbiome composition that improves feed digestion and utilization, while precluding presence of harmful microbes or composition thereof.

## DECLARATIONS

### Ethics approval and consent to participate

It was not considered to ask for an authorization because the procedures used in animals were those used under a common clinical veterinary procedure, therefore not subject to regulation by the Spanish and European Legislation related with the protection of animals used for scientific purposes. Nevertheless, the animals were manipulated according to the Spanish Policy for Animal Protection RD 53/2013, which meets the European Union Directive 86/609 about the protection of animals used in experimentation.

### Consent for publication

Not applicable

### Availability of data and material

The datasets during and/or analysed during the current study available from the corresponding author on reasonable request, and the authors plan to upload them to a data repository soon.

## Authors’ contributions

OGR performed the analyses of sequence data, managed the genotyping of the animals, implemented the statistical analyses, discuss the results and wrote the first draft of the manuscript. AGR, IZU and RAT made the experimental design and executed the experiments, collected and analyzed the samples, discussed the results and helped to write the manuscript. AHU performed the preparatory actions for the sequencing analysis and helped writing the manuscript. All authors read and approved the final manuscript.

## Acknowledgments

Authors wish to thank Iurancha González for assistance during sampling and Beatriz Oporto and Medelin Ocejo for DNA preparations, as well as the staff from Escuela Agraria de Fraisoro (Basque Department of Environment, Rural Development, Agriculture and Fishery).

---

**Figure S1.**
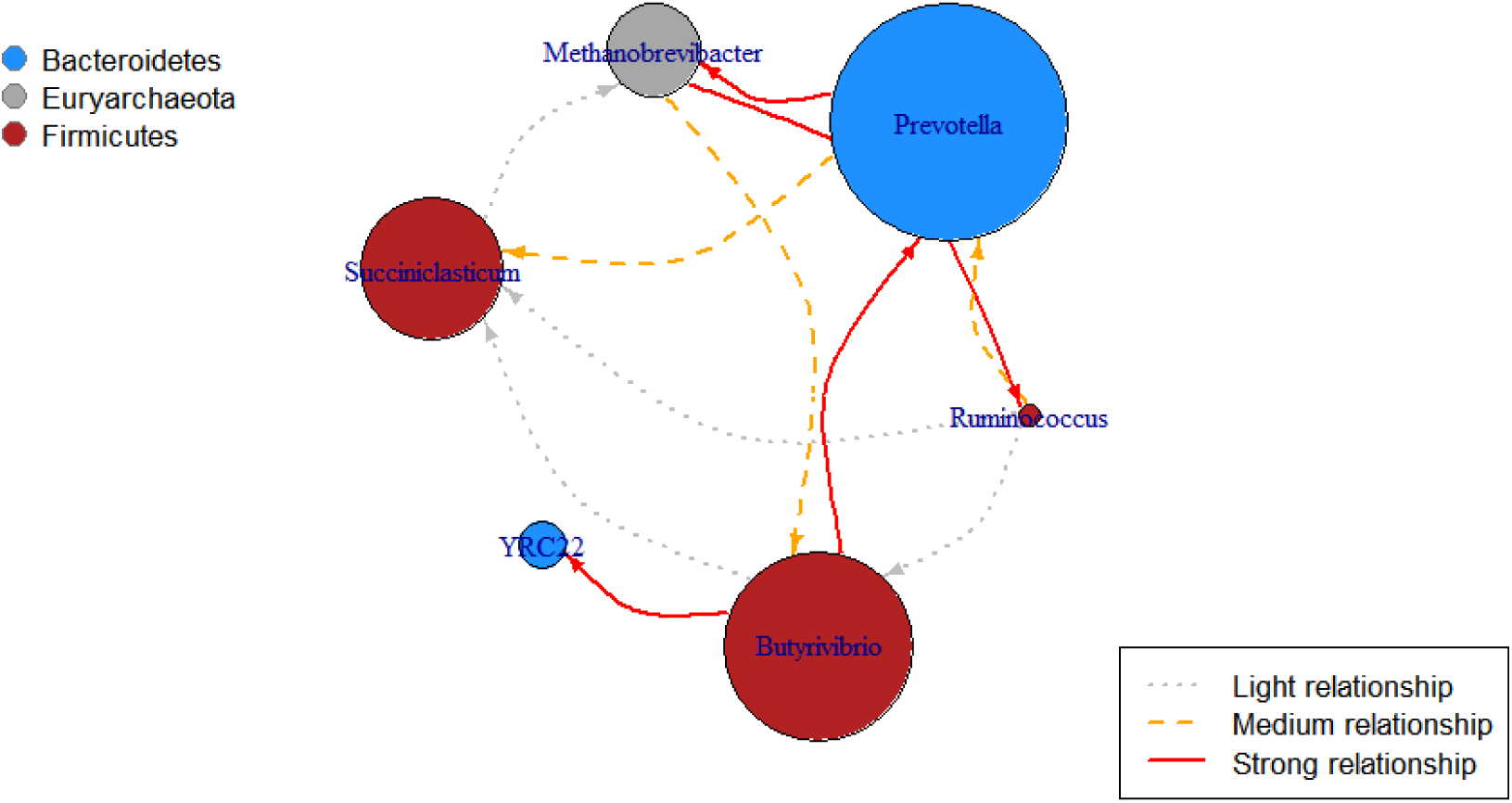
Microbial network based on 16S rRNA-gene based region for microorganism with relative abundance larger than 0.1% in both breeds. The size of the nodes represents the relative abundance of the genera.

**Figure S2.**
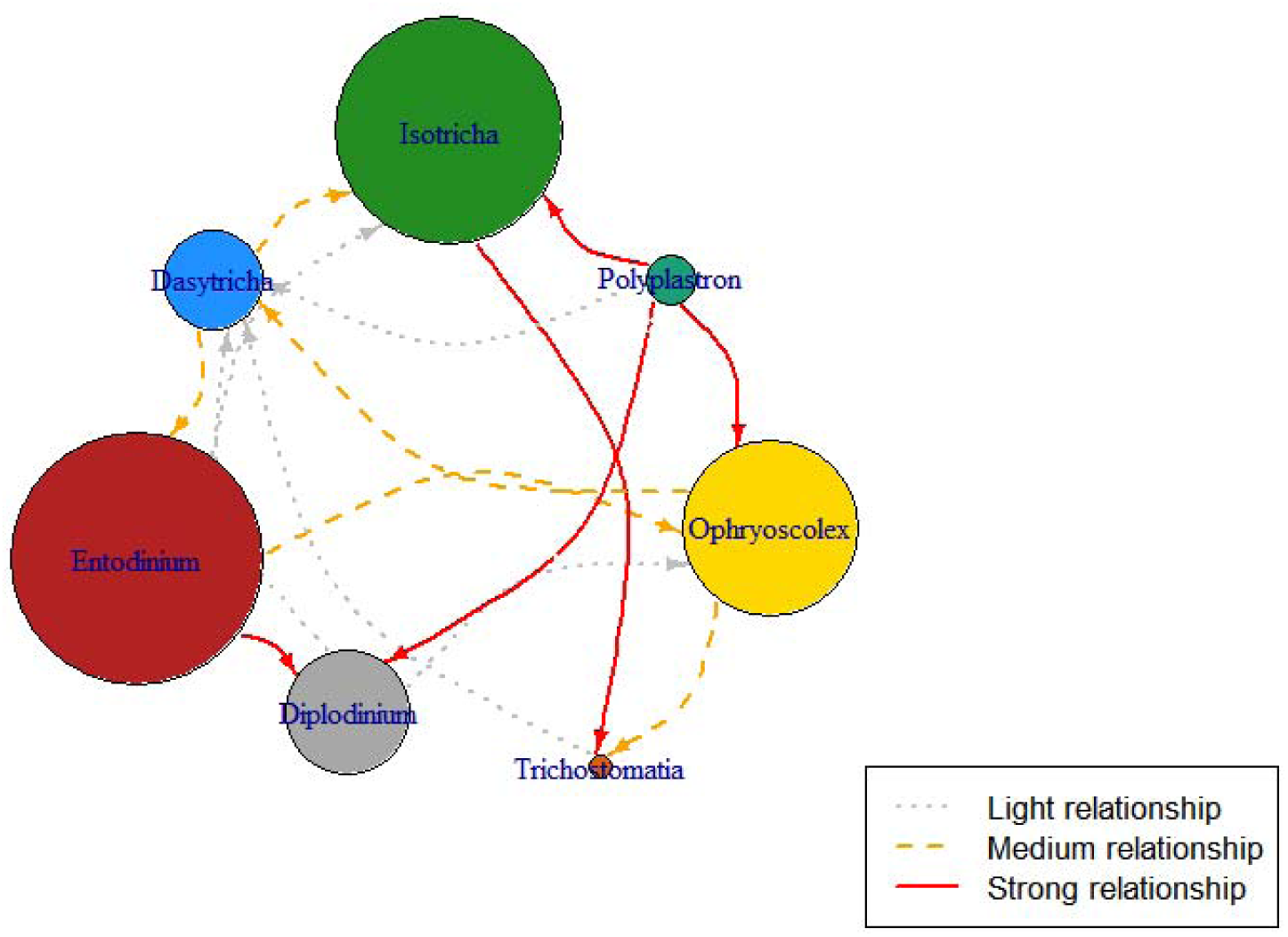
Microbial network based on 18S rRNA-gene based region for ciliates with relative abundance larger than 0.1% in both breeds. The size of the nodes represents the relative abundance of the genera.

## Abbreviations

FDR: False Discovery Rate
NGS: Next Generation Sequencing
MAF: Minor Allele Frequency
OTU: operational taxonomic units
PC: Principal components
RA: Relative abundance
SNP: Single Nucleotide Polymorphisms

